# xRead: a coverage-guided approach for scalable construction of read overlapping graph

**DOI:** 10.1101/2023.05.23.541864

**Authors:** Tangchao Kong, Bo Liu, Yadong Wang

## Abstract

The development of long-read sequencing is promising to high-quality and comprehensive de novo assembly for various species around the world. However, it is still challenging for genome assemblers to well-handle thousands of genomes, tens of gigabase level genome sizes and terabase level datasets simultaneously and efficiently, which is a bottleneck to large de novo sequencing studies. A major cause is the read overlapping graph construction that state-of-the-art tools usually have to cost terabyte-level RAM space and tens of days for that of large genomes. Such lower performance and scalability are not suited to handle the numerous samples to be sequenced. Herein, we propose xRead, an iterative overlapping graph approach that achieves high performance, scalability and yield simultaneously. Under the guidance of its novel read coverage-based model, xRead uses heuristic alignment skeleton approach to implement incremental graph construction with highly controllable RAM space and faster speed. For example, it enables to process the 1.28 Tb *A. mexicanum* dataset with less than 64GB RAM and obviously lower time-cost. Moreover, the benchmarks on the datasets from various-sized genomes suggest that it achieves higher accuracy in overlap detection without loss of sensitivity which also guarantees the quality of the produced graphs. Overall, xRead is suited to handle numbers of datasets from large genomes, especially with limited computational resources, which may play important roles in many de novo sequencing studies.

## Background

De novo assembly is to reconstruct the sequence of donor genome from reads without reference which is fundamental to genomics studies. The rapid advances in long-read sequencing technologies, such as Single Molecule Real Time (SMRT) sequencing [1] and nanopore sequencing [2], have been able to produce reads having >10kbp median length and >100kbp maximum length [3]. They have superior repeat-spanning ability to resolve complex repetitive regions which greatly helps to achieve high-quality assemblies such as telomere-to-telomere assembly [4] and haplotype assembly [5]. However, the assembly of large genomes with tens of gigabases length (such as *P. taeda* [6], *A. mexicanum* [7], *E. superba* [8], etc.) is still non-trivial. One of the bottlenecks is the computation-intensity, i.e., most of the state-of-the-art assemblers have to cost thousands of CPU hours and require terabytes of RAM space for such tasks [9, 10]. In this situation, employed tools could be not scalable to handle many large genomes with commonly used computational environments, since the real-time cost could be prohibitive and the availability of computers having large RAM configurations could be also limited. Thus, there could be still technical bottleneck to large-scale de novo sequencing studies like Vertebrate Genomes Project [11] and Earth Biogenome Project [12].

A primary cause of the bottleneck is the alignment of reads, since most of the long-read assemblers are in Overlap-Layout-Consensus (OLC) approach which needs all-against-all read alignment to construct an overlapping graph at first [13-18]. This initial step has quadruplicate time complexity in theory, while its RAM space cost is also high if all the read information is kept in memory. Moreover, the combination of complex genome repeats and high sequencing errors also make it a difficult task to implement high-quality graph construction. State-of-the-art tools use various heuristics to accelerate the speed, reduce RAM cost and improve the accuracy and sensitivity of read overlapping.

Seed-and-extension is one of the most commonly used heuristics. Most of such approaches retrieve short matches (i.e., seeds) between various reads (usually through some specifically designed index data structures) and then implement extended alignments around them to confirm the actual overlapped parts of the reads. HGAP [19] is one of the earliest tools tailored to the assembly of noisy long reads. It heuristically indexes a proportion of the longest read and employs a typical seed-and-extension alignment tool (BLASR [20]) to align them with other reads. FALCON assembler [14] employs DALIGNER [21] which partitions reads into blocks and uses sorted k-mers within them as the index, further, the blocks are merged to discover read overlaps. Wtdbg2 [17] indexes a quarter of k-mers as seeds and takes each tiling 256 bp subsequence as a bin for each read. Further, it also employs 256 bp-bin-based dynamic programming for extension instead of base-level alignment. Flye [22] collects frequent k-mers in reads as seeds and estimates the overlaps by finding the longest common sub-path with a fast dynamic programming algorithm. Shasta [18] uses randomly selects k-mers (seeds), finds candidate overlaps with the LowHash algorithm, and performs a tailored marker alignment approach for extension. Minimap2 [23] is a generic aligner suited to find the overlaps of long reads in various lengths and error rates which is also employed by several state-of-the-art assemblers such as Raven[24], PECAT[25], and Nextdenovo[8]. It essentially uses minimizer-based seeding and chaining to detect read overlaps and also supports the base-level alignment of anchored reads if necessary.

Some of the previous studies also focus on the acceleration of the base-level alignment which is helpful to reduce the time cost of extension step. Most of them take advantage of Single Instruction Multiple Data (SIMD) instructions such as Intel AVX instructions or Compute Unified Device Architecture (CUDA) in Nvidia GPU. Manavski and Valle proposed an implementation of Smith-Waterman algorithm under CUDA framework. Libssa [27] uses AVX2 instructions to accelerate the classical Smith-Waterman and Needleman-Wunsch algorithms. Parasail [28] is a SIMD-based implementation of global, semi-global and local alignments which supports a couple of instruction sets such as SSE2, SSE4.1, AVX2, AltiVec and NEON. Suzuki and Kasahara developed a fast SIMD-based alignment algorithm named libgaba [29] and it was further improved in KSW2 [23] and employed by Minimap2.

Although many efforts have been made, the overall cost of read overlapping is still high under the seed-and-extension framework due to many issues such as large the number of reads, the high sequencing errors, the ubiquitous repeats in donor genomes, etc. Alignment-free approaches are also developed and employed by state-of-the-art assemblers. Most of them use tailored compact sequence representations, i.e., sketches [30], to directly measure the similarities between reads. MHAP [13] uses MinHash technique which employs 256-1512 hash functions to construct sketches and use them to estimate the Jaccard similarity of the reads. Canu [15] uses adaptive k-mer weighting to improve MinHash which reduces the effect of highly repetitive k-mers. MECAT [16] splits reads into blocks and finds candidate overlaps with at least one matched block. Low-similarity overlaps are then filtered by removing k-mers with low distance difference factor (DDF) scores. NECAT [31] extends DDF scoring by sorting all k-mer pairs and chaining them together to remove false positive k-mers which is more suited to the heterogeneous sequencing errors of ONT reads. Such alignment-free approaches avoid the computationally intensive base-level alignment, however, their time cost is still non-neglectable since it usually needs many query and merging operations to make a number of sketches to handle sequencing errors and achieve sensitivity.

Long-read assembly approaches with higher scalability and speed are in wide demand to deal with the ever-increasing sizes and numbers of de novo sequencing genomes. Moreover, there are still a number of false positives/negatives in overlapping graphs caused by the various tradeoffs of existing tools on sensitivity, accuracy and performance. Herein, we proposed xRead, an incremental overlapping graph construction approach that is able to achieve high scalability, performance and yields simultaneously. Guided by a novel read-coverage-based objective function, xRead iteratively builds and refines the overlapping graph with heuristic read indexing and lightweight alignment skeletons. The approach has three major contributions. Firstly, it has outstanding scalability for memory usage which enables to build the overlapping graphs of high-coverage sequencing datasets of very large genomes with low and tunable RAM space cost, e.g., building an overlapping graph of 32X PacBio sequencing dataset (1.9 Terabyte) of Axolotl genome with 64GB or lower RAM. Secondly, it has high speed for various-sized genomes, e.g., several times faster on average than that of Minimap2 on the long-read datasets from *E. coli* to human genomes. Thirdly, it enables to produce highly accurate read overlaps without loss of sensitivity, which is helpful to high-quality graph construction. xRead is suited to handle many ONT and PacBio datasets from large genomes effectively and efficiently in de novo sequencing studies.

## Results

### Overview of the xRead approach

xRead is motivated by a simple assumption that the read depth is homogeneous along the sequenced genome. This is essentially the case due to the less bias of long-read sequencing [32-34]. As the real sequencing coverage is unknown, xRead builds the overlapping graph under the guidance of aligned read coverage as it indirectly reflects the sequencing coverages of all the local genomics sites. The overlapping graph is iteratively built (a schematic illustration is in Fig. 1). In each iteration, xRead implements graph construction and refinement in three major steps as follows.

**Fig 1.**
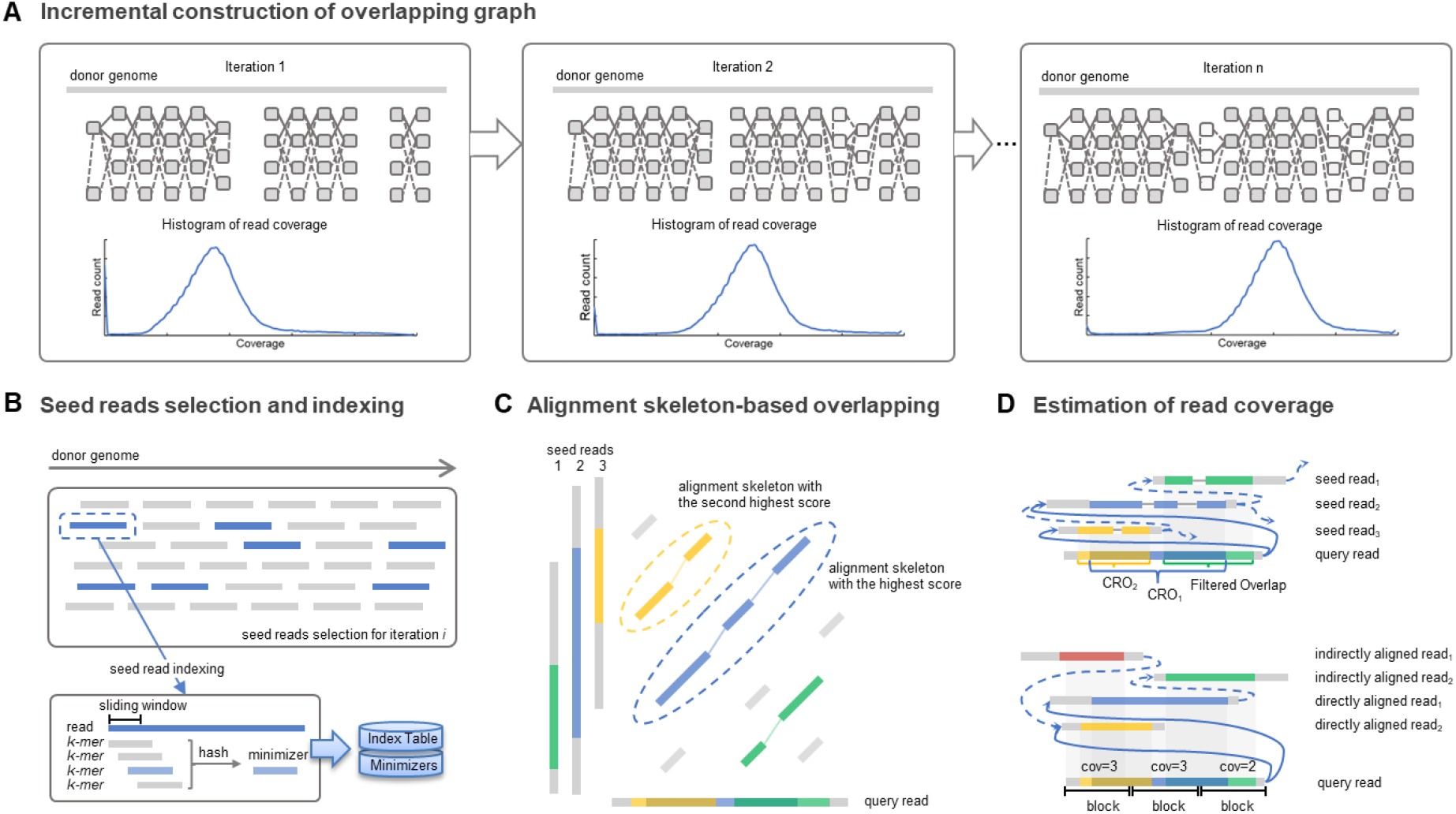
A schematic illustration of xRead. **a**. The incremental construction of overlapping graph. The subplots represent the connections among reads being incrementally recovered within various iterations. The gray blocks indicate the sequencing reads. The dashed and solid lines respectively indicate the overlaps between the seed and the query reads, and the seed reads themselves. The histograms in the lower part indicate the distributions of read coverage which are updated in various iterations. **b**. The selection and indexing of seed reads. xRead selects a portion of lowly covered reads as seed reads (marked as blue bars) and a list of minimizers is extracted using a hash function. A minimizer-based index is then built by a hash table-based data structure. Meanwhile, the same hash function is also used to generate minimizers for query reads. **c**. Alignment skeleton-based read overlapping. For a given query read, xRead finds the MBs (marked as colored bars) between it and all the seed reads via the index. Further, it uses the SDP approach to generate one or more alignment skeletons (the dashed ovals indicate the skeletons with the first and second highest scores). **d**. Estimation of read coverage. The upper subplot represents the selection of CROs. The first two highest-scored skeletons are selected as CROs which bring new edges (represented by solid lines) to the overlapping graph. Other skeletons with lower scores are filtered out. The lower subplot represents the (re-)estimation of read coverage. xRead splits the reads into non-overlapping blocks and then counts the reads directly (marked as solid lines) and indirectly (marked as dashed lines) being aligned to it.

1) xRead selects a proportion of reads having relatively low coverages and high lengths as seed reads and builds a partial read index for them.

2) xRead employs a lightweight alignment skeleton approach to discover new read overlaps between the seed reads and other less covered reads (termed as query reads).

3) xRead constructs and refines the overlapping graph based on the produced alignment skeletons. Further, it estimates the read coverages and ends the process if there is no read having low enough coverage, otherwise, turns to step 1 for a new iteration.

This design is tailored to achieve a highly scalable graph construction since the RAM cost can be well-handled with the proportion of the reads being indexed. Moreover, the time cost is also relatively low with the lightweight alignment skeleton approach while it also has the ability to produce accurate read overlaps similar to classical base-level alignment. Refer to Methods section for more details about the implementation.

### Benchmarks on simulated datasets

We implemented benchmarks on simulated long-read datasets from nine genomes whose sizes are from several megabases to several tens of gigabases (Supplementary Table 1) to assess the baseline performance of xRead. A 50x coverage ONT-like dataset was simulated (Supplementary Table 2) for each of the nine genomes by PBSIM2 with its pre-trained R103 chemistry model and recommended error ratio (substitution: insertion: deletion=23: 31: 46). The mean read length and total error rates were configured as 13kbp and 13%, respectively, referring to the previous study [3]. The generated MAF files were used as ground truth. It is also worth noting that we divided the chromosomes of *A. mexicanum* genome into at most 1Gbp segments before simulation due to the limit of PBSIM2 on chromosome length. The runtime, memory footprint, sensitivity and accuracy of various tools are in Fig. 2 and Supplementary Tables 3-4. Four state-of-the-art read overlapping tools, MHAP, MECAT2, Minimap2, and wtdbg2 were also employed for comparison (refer to Methods section for more detailed information). Mainly, three issues were observed from the results as follows.

**Fig 2.**
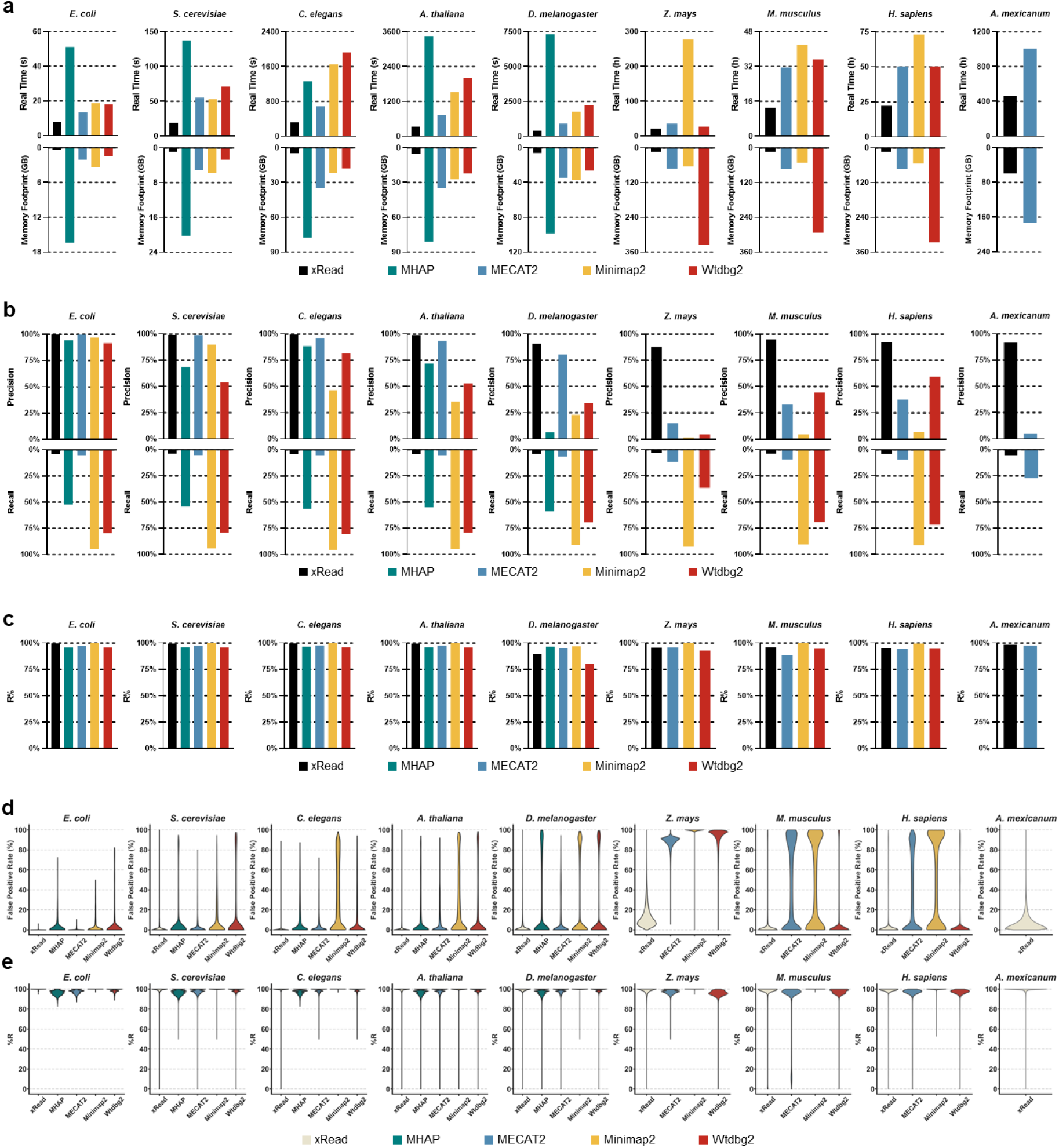
Results on simulated datasets. The figure depicts the real-time, peak memory **(a)**, precision, sensitivity **(b)**, R% **(c)**, and violin plots of FPR **(d)** and R% **(e)** of each overlapper on nine simulated datasets of various-sized genomes (from *E. coli* to *A. mexicanum*). **a-c**. In each subplot of **(a), (b)**, and **(c)**, the color black, green, blue, orange, and red respectively represent the overlapper xRead, MHAP, MECAT2, Minimap2, and wtdbg2. **d-e**. The violin plot of FPR and R% for various datasets. For a given tool, the absence of the results for some datasets is due to its failure during benchmarking. Refer to supplementary material for detailed information of datasets and results.

Firstly, xRead has outstanding scalability for genomes of various sizes.

xRead allows configurable RAM usage, and we limited its RAM space to 16GB, 24GB and 64GB during benchmarking for the datasets of small (*E. coli, S. cerevisiae, C. elegans, A. thaliana*, and *D. melanogaster*), large (*Z. mays, M. musculus* and *H. sapiens*) and very large (*A. mexicanum*) genomes. A larger RAM space is used for *A. mexicanum* to reduce some I/O operations although it is still obviously lower than that of the other four tools. xRead successfully accomplishes all the tasks (Fig. 2a and Supplementary Table 3), suggesting that the read coverage-guided iterative design is able to well-handle large sequencing datasets with medium size workstations or servers. Meanwhile, the speed of xRead is about 1.2-18 times faster (in real-time) than that of other tools for the datasets. Overall, the performance suggests that xRead has good scalability to various-sized genomes.

Other tools were run without any limit on memory during the benchmark and their memory footprints are 3-47 times higher than that of xRead with relatively lower speed. This could be caused by the two kinds of employed read overlapping strategies as follows. One is to load all the reads and process them in memory, like MHAP and wtdbg2, whose RAM usage is quite high. The other one is to divide the whole datasets into batches of reads and separately handle each of them, like MECAT2 and Minimap2. For each batch, the involved reads are indexed in time and other reads are aligned to them to discover overlaps, which is more similar to xRead. However, they also required significantly higher RAM space, e.g., MECAT2 used over 172GB RAM to process the *A. mexicanum* dataset with 64 CPU threads. Moreover, the lower speed of MECAT2 and Minimap2 could be also due to their different designs, i.e., xRead adaptively selects a set of reads having fewer overlaps to index and align while MECAT2 and Minimap2 straightforwardly divide the datasets and align all the reads to a specific batch of reads in each iteration. It is also worth noting that only xRead and MECAT2 finished the *A. mexicanum* task, and the real-time of xRead is less than 40% of that of MECAT2 which saved about 25 days. On this dataset, MHAP and wtdbg2 ran out of memory (over 1Terabytes). For Minimap2, it showed relatively low speed, i.e., it cost about 178 hours (with 64 CPU threads) to process only 11 read batches (about 3.29% of the dataset). Considering the high time cost (estimated over 225 days), we early stopped the program.

Secondly, xRead is able to produce accurate read overlaps without loss of sensitivity.

We evaluated the overall precisions and sensitivities of the tools (Fig. 2b and Supplementary Table 4, Precision and Sensitivity columns). Since too short overlaps could be caused by coincidence and false positives which do not make sense to genome assembly, only the read overlaps longer than 500bp were considered in the evaluation referring to previous studies [13]. The results of the tools varied, indicating the different tradeoffs in their designs. With the relatively conservative CRO strategy, xRead output a set of “core overlaps” which connect each of the seed reads and all the other reads with high scores, and it achieved the highest precisions on all the datasets. MHAP, MECAT2, and wtdbg2 are more likely to pursue the balance between accuracy and sensitivity, so that their sensitivities are higher than that of xRead but precisions are lowered. Minimap2 tries to detect read overlaps comprehensively. It outputs the highest number of read overlaps and achieved the highest overall sensitivities on all the datasets. However, its precision is lowest among the tools, i.e., there are also high numbers of false positives in the produced graphs for not only the large but also the relatively small genomes, such as *A. thaliana* and *D. melanogaster*, which could affect the layout of the reads during genome assembly.

We further investigated more detailed information of the graphs produced by the tools with three additional three metrics, R%, C% and Con. Num. The results indicate that xRead is still without loss of sensitivity and has the potential to support high-quality assembly, although its outputs only covered a proportion of ground truth overlaps.

R% indicates the proportion of the reads having at least one ground truth overlap being recovered. The result (Fig. 2c and Supplementary Table 4, R% column) suggests that Minimap2 has the highest R% due to its sensitivity. For other tools, their R% are quite close to each other, moreover, in absolute terms they are also very high and comparable to that of Minimap2, i.e., most of the reads have one or more correct overlaps being recovered. For xRead, this indicates that most of the reads can be either directly aligned or indirectly connected via seed reads and only a very small proportion of the reads had no correct connection at all in the graphs. Therefore, it provides the opportunity to more sensitively recover read overlaps to produce comprehensive graphs (see more results below).

C% indicates the percentage of the donor genome being covered by the connected reads. Herein, connected reads indicate the reads having at least one edge (overlap) in the produced graph and this metric measures the coverage of the graph on the donor genome. It was observed that for various datasets all the produced graphs had very high and close C% (nearly 100%) with non- or very few gaps (i.e., uncovered genomic regions, the C% and Gap Num. columns of Supplementary Table 4). We further found that the <100% C% were mostly caused by the simulated reads that did not fully cover the genome, not the fault of the overlapping tools. For xRead, this result suggests that the seed- and their connected reads can cover the whole genome. Thus it is helpful for assemblers to fully recover the sequence of the donor genome, since no local region is missed during graph construction.

Con. Num. indicates the number of connected components of produced graphs which measures their connectedness. It was observed (Con. Num. column of Supplementary Table 4) that on various datasets nearly all the produced graphs had relatively small numbers of connected components. This is partially due to the superior read lengths, meanwhile, it also indicates that tools enable the effective alignment and connection of the read. With the highly connected graphs, all the tools have the chance to indirectly infer the read overlaps in the same components via transitive relationships and achieve high sensitivity. For xRead, the numbers of connected components are comparable to that of other tools on various datasets, however, it slightly increased for some of the genomes. We investigated the graphs and found that most of the reads were connected and clustered in a few components, and other components only had very few (e.g., two or three) reads. This is mainly due to that a small proportion of selected seed reads are error-prone, some of the non-seed reads were aligned to them by accident. It usually happened in large datasets, since more outlier reads with serious sequencing errors occurred with higher total read numbers. Other tools also had such a trend, moreover, by filtering out those extremely small ones, all the tools had very similar numbers of connected components.

Considering the precision, coverage and connectedness, we realized that the graphs produced by xRead are suited to function as a core graph to guide successful read assembly or error correction, referring to previous studies [19]. Furthermore, xRead also provides an additional tool to optionally expand the core graphs to comprehensive graphs which is suited to meet the various requirements of genome assembly approaches. Mainly, it is implemented by a transitive rule-based width-first searching approach to iteratively recover the overlaps among non-seed reads (refer to Methods section for more detailed information). The tool enables to recover as many read overlaps as possible to achieve high sensitivity, meanwhile, it also supports to fine-tune the number of iterations for various tradeoffs between sensitivity and precision. The overall sensitivities of the graphs expanded by 1, 3, and 5 iterations are in Supplementary Table 5. It is observed that the sensitivity of the produced graphs greatly improved by implementing even only one iteration expansion. Moreover, the number of overlaps saturated after 5 iterations for most of the reads and the sensitivity metrics (overall sensitivity and R%) are higher or close to that of Minimap2.

Thirdly, xRead is able to keep high yields along the whole genomes.

We assessed the read overlaps in various local regions to investigate the behaviors of xRead under various genomic contexts. Mainly, the reference genome was split into non-overlapping blocks (size: 1kbp and 10kbp for small and large/very large genomes, respectively) and the reads were assigned to corresponding blocks according to the ground truth. Further, the false positive overlap rates (FPR) and R% in various blocks were separately calculated. Herein, for a given block, FPR is defined as the proportion of false positives against all of the overlaps that at least one of the involved reads belongs to the block, and R% is the proportion of the reads belonging to the block but having no ground truth overlap recalled. The violin plots of FPR and R% for various datasets are in Fig. 2d and e.

The FPR plots show that xRead is able to keep relatively low FPR along the whole genome and has a significantly higher number of blocks with zero FPR. We further investigated the high FPR blocks of xRead and found that most of them are of complex repetitive regions. Such cases are quite complicated and challenging due to the combination of the high similarity of repeat copies and the serious sequencing errors. An example is the *Z. mays* dataset that the genome is highly repetitive and all of the tools cannot maintain very low FPR. However, with the relatively conservative CRO strategy, xRead still maintained relatively high precision, i.e., for most of the blocks it had <20% FPR while other tools produced ubiquitous high FPR (>80%) blocks.

The R% plots show that all of the tools have high R% along the whole genome, i.e., they have the ability to discover real overlaps for the reads in various regions. For xRead, this derives from the nearly non-gap coverage of the whole genome by the seed reads and the high ability of alignment skeletons to capture the non-seed reads from the same local regions, which also depicts the relatively high quality of the core graph. We further investigated the detailed information of the low R% blocks of xRead and observed that they also concentrated in highly repetitive regions. This is mainly due to that the reads from such regions, especially the ones having relatively short lengths and lower repeat spanning ability, are more likely to be aligned to the seed reads from other copies of the same repeats and the incorrect CROs mislead the calculation of read coverage. Thus, such reads achieved high coverage in the first several iterations of graph construction and had no chance to be rescued later.

### Benchmarks on real datasets

The tools were further benchmarked with seven real sequencing datasets from ONT and PacBio platforms (Supplementary Table 6). Three are from relatively small genomes, i.e., *E. coli, C. elegans* and *D. melanogaster*, and produced by ONT platforms. Three are human datasets from GIAB sample HG002 (NA24385), two of them are ONT datasets in fast and super high accuracy base-calling modes, respectively, and the other one is PacBio HiFi dataset. The ONT super high accuracy and PacBio HiFi datasets were employed to assess the ability of xRead on high-quality long-read datasets. A PacBio CLR dataset from *A. mexicanum* genome was employed to assess the tools on very large genomes.

Due to the absence of ground truth, we take advantage of the high mappability of long reads to produce pseudo ground truth. That is, for each dataset, the reads were aligned to the corresponding reference genome using Minimap2 with default settings. The reads being unaligned or in low mapping quality were marked as ambiguous reads and filtered out. The remaining reads as well as their mapping positions were used to compose the pseudo ground truth overlap set. The performance, accuracy, and sensitivity of the tools were then assessed. The same limitation on RAM space was still applied for xRead and no limitation for other tools during benchmarking.

The results on the real datasets showed similar trends to that of simulated datasets. xRead still had high performance (Fig. 3a and Supplementary Table 7), i.e., overall faster speed with lower memory footprints than that of other tools. It is also worth noting that xRead also had higher performance on the ONT super high accuracy and PacBio HiFi datasets (several to tens of times faster than other tools), indicating that it is not only suited to noisy but also high-quality long reads. An exception occurred on the ONT fast mode human dataset that the speed of xRead is slower than that of MECAT2 but still comparable. It is mainly caused by the repeats of the human genome, i.e., xRead spent much time on the sorting and linking of the numerous minimizers from highly repetitive regions. Although higher, the time cost is still acceptable in absolute terms. The performance also shows that xRead is more suited to handle very large genomes. It is the only tool that accomplished the real *A. mexicanum* dataset successfully on the employed server. MHAP and wtdbg2 were out of memory (>1 Terabytes). MECAT2 raised an error signal and terminated. The speed of Minimap2 was still very low like that of the simulated datasets and was early-stopped considering its very high estimated time cost.

**Fig 3.**
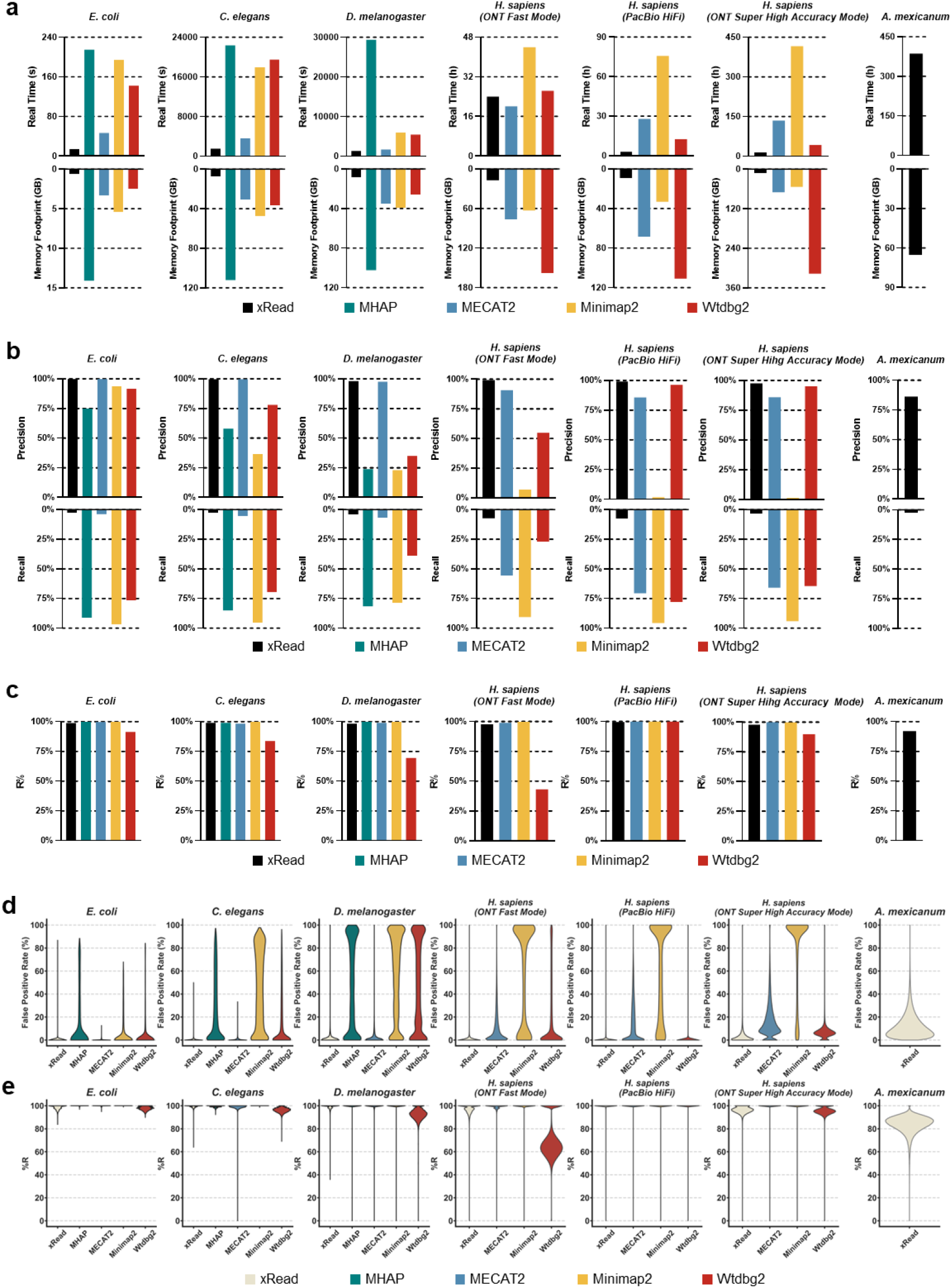
Results on real datasets. The figure depicts the real-time, peak memory **(a)**, precision, sensitivity **(b)**, R% **(c)**, and violin plots of FPR **(d)** and R% **(e)** of each overlapper on seven real datasets of various-sized genomes (*E. coli, C. elegans, D. melanogaster*, three *H. sapiens* dataset from different platforms, and *A. mexicanum*). **a-c**. In each subplot of **(a), (b)**, and **(c)**, the color black, green, blue, orange, and red respectively represent the overlapper xRead, MHAP, MECAT2, Minimap2, and wtdbg2. d-e. The violin plot of FPR and R% for various datasets. For a given tool, the absence of the results for some datasets is due to its failure during benchmarking. Refer to supplementary material for detailed information of datasets and results.

The precision of xRead was still higher than that of other tools on the real datasets (Fig. 3b and Supplementary Table 8, Precision column). MECAT2 also achieved similar precision on smaller genomes, however, it decreased on the three human datasets, potentially due to the characteristics of its DDF score model. The precision of wtdbg2 was relatively high on PacBio datasets while significantly lowered on ONT datasets, indicating that it could be not suited to handle ONT reads. The precisions of MHAP and Minimap2 were significantly lower on all the datasets.

The overall sensitivities of the tools still varied due to their different designs, while Minimap2 was the highest one in contrast to its precision. On the R% metric (Fig. 3c and Supplementary Table 8, R% column), the tools were high and close to each other (except for wtdbg2 on ONT datasets), indicating that they also had to-some-extent similar abilities to detect the overlaps of real sequencing reads. Mainly, the reads having no true positive overlap recalled were from highly repetitive regions. In this situation, all of the tools were likely to be affected by the reads from various copies of the repeats and were hard to detect overlaps correctly.

In addition, we also found another two issues which slightly lowered the R% of xRead. One is the interaction of the repeats in genomes and the serious sequencing errors in some parts of the reads (a more detailed discussion is given below). The other is the lack of ground truth. That is, most of the ambiguous reads are potentially from the incomplete regions (i.e., the regions marked as “N” bases) of reference genome and cannot be mapped. The overlaps with such reads were discarded in the evaluation since their correctness cannot be determined. Thus, some of the reads were recognized as non-overlapped as the ambiguous reads were the seed reads they overlapped with. We checked a portion of such overlaps and realized that they were also reasonable. That is, most of them were from ambiguous reads to the reads that can be mapped nearby incomplete regions with high scores. Supplementary Table 9 gives more detailed information about the proportions of the reads having no correct overlap caused by various issues. Overall, considering that the overall R% of xRead is still high in absolute terms, it could be not problematic for genome assembly.

The metrics about the coverage and connectedness of the produced graphs were further assessed (C%, Gap Num. and Con. Num. columns of Supplementary Table 8). Similarly, the graphs are highly connected (i.e., relatively low numbers of connected components) and able to cover nearly the whole genomes with few gaps. It is also worth noting that for all of the tools the number of connected components slightly increased. This is also due to that real sequencing reads are more error-prone and more likely to cause small components to have a few accidentally aligned reads. A partial evidence is that all the tools have obviously lower numbers of connected components on the human PacBio HiFi dataset than that of the ONT fast mode dataset, putatively due to the higher sequencing quality. However, such small and noisy components could also not seriously affect genome assembly since it is not difficult to implement filtration by the size and quality of the overlaps. Furthermore, we also implemented graph expansion for the graphs produced by xRead (Supplementary Table 10). Consistent with that of simulated datasets, i.e., the overall sensitivity of the graphs obviously improved and nearly saturated after 5 iterations.

The read overlaps in local regions were also investigated. The violin plots of FPRs and R% of various genomic blocks are in Fig. 3d and e. xRead had the highest number of blocks with zero FPR and the distributions of the R% of various tools were quite similar. Moreover, it was observed that most of the reads having false positive and missing overlaps concentrated in a few blocks, leading to local high FPRs and low R%. We further investigated those blocks and found that the false positives mainly derived from highly repetitive regions. However, by separately investigating low R% blocks, we observed that the overlaps missed by xRead were caused by more complicated interactions of sequencing errors and repeats as follows.

Firstly, most of the missing overlaps were caused by error-prone read parts. In real data, some parts of the long reads have very high sequencing errors, so there is a lack of matches in those local regions. If the error-prone part is very long and in the inner region of the read, a lowly scored skeleton could be produced and mistakenly filtered out by xRead. Another case is as mentioned above that the low-quality part is nearby either of the ends of the read where xRead could skip the skeleton due to the placement of the overlap.

Secondly, a small proportion of the missing overlaps were due to repeats as well as the design of xRead. During the construction of alignment skeletons, xRead initially does not use highly repetitive minimizers which occur more than a threshold of times in the index since most of them bring false positive matches which may affect both precision and performance. In most cases, the remaining minimizers are enough to build correct alignment skeletons. However, some of the reads from repetitive regions could lack matches and xRead employs an additional approach to solve the problem. That is, for the low-scored skeletons having a high number of repetitive minimizers, xRead uses the initial matches of the less repetitive minimizers as anchors and re-searches the matches between repetitive minimizers around the anchors to refine the skeletons. This helps to resolve most of the reads from repetitive regions. However, for only a very small portion of the reads, they have relatively short lengths and are fully enclosed in long repeats. Their skeletons are of low specificity and could bring many false positives so that xRead discards them.

## Discussion

Long-read sequencing technologies are promising for the high-quality genome assembly of various species. However, it also makes new requests to the scalability of de novo assembly tools to handle many hundreds to thousands of genomes, tens of gigabase level genome sizes and terabase level datasets simultaneously and efficiently. As one of the most computationally-intensive steps, the construction of the read overlapping graph gets challenged, especially with limited computational resources. It is in wide demand to develop novel algorithms and tools having high scalability and performance. Herein, we propose xRead, a novel approach tailored for scalable overlapping graph construction. The benchmarks on simulated and real datasets suggest that xRead is able to well-handle various-sized genomes and long-read datasets with controllable RAM usage and its performance is higher than that of state-of-the-art tools. It is also able to achieve high precision without loss of sensitivity as well. Mainly, xRead has three technical features as follows.

Firstly, the read coverage-guided model not only enables the iterative construction of an overlapping graph with partial read information, but also adaptively checks the completeness of the graph in construction and focuses on the reads being less overlapped. This feature realizes the controllable RAM usage which is critical to achieve high scalability. Moreover, it also prunes a large number of unnecessary read alignment operations to improve the performance from high-level design.

Secondly, the minimizer-based alignment skeleton has similar ability to detect read overlaps as full alignment, but its speed is much faster. Moreover, with the flexible connections between local matches, this approach also has a good ability to deal with ubiquitous sequencing errors.

Thirdly, the skeleton scoring and selection system not only considers the sequence similarity but also the placement of the aligned read parts, furthermore, it conservatively selects only the most likely candidate overlaps. This strategy helps to reduce false positives. Moreover, it also keeps the sensitivity to detect the overlaps between seed and non-seed reads which enables xRead to achieve high graph coverage on donor genome and high connectedness which are helpful to downstream analysis.

Although it achieves overall scalability, performance and yields, xRead could still be affected by sequencing errors and genome repeats like that of other state-of-the-art tools. Serious sequencing errors may lead to missing overlaps due to the lack of matches. However, with the improving sequencing quality of ONT and PacBio platforms, the number of highly error-prone reads greatly decreases. Besides that, such error-prone reads could also bring uncertainty in downstream steps so that straightforwardly filtering out the non-overlapped reads could be an acceptable option as well. Repeats are of intrinsic genomic property and could mislead xRead to produce either false positives or false negatives. Essentially, the problem derives from the cases in which relatively short reads are fully enclosed in long repetitive regions. Such reads have very low mappability which is hard to solve in theory. However, this is also the case for only a very small proportion of reads in real ONT and PacBio data and it could be not very problematic for genome assembly. Moreover, the problem would be further mitigated by the ever-increasing read length. However, the crosstalk of serious sequencing errors and complex genome repeats is still an open problem to not only the initial overlapping graph construction but also nearly all the steps of de novo assembly. And it is also an important future work for us.

On a higher level of view, it is important to achieve sensitivity and accuracy simultaneously, however, there are always tradeoffs in the practical implementations of various read overlapping tools. Unlike many existing tools which are to detect as many as possible overlaps, xRead is to some extent in a minimalist design that constructs a simplified, but not over-simplified, overlapping graph through the selection and alignment of seed reads. This design is beneficial to implement highly scalable and efficient graph construction even if for very large donor genomes and huge numbers of sequencing reads, which breakthroughs the bottleneck to large-scale de novo assembly tasks. Moreover, benchmarks show that the high precision, coverage, and connectedness enable the produced graph to function as a core graph for downstream steps such as read layout and error correction. Further, it also has the potential to recover read overlaps more comprehensively to meet the various requirements of assembly tools via transitive rule-based inference.

## Conclusion

The construction of the overlapping graph is fundamental to long read-based genome assembly and state-of-the-art tools could have bottlenecks to handle the ever-increasing sizes of genomes and datasets in many ongoing and upcoming large-scale de novo sequencing tasks. xRead takes advantage of a read coverage-based iterative construction approach to simultaneously achieve high scalability, performance and yields. We believe that it may lay a new foundation for long read-based genome assembly and play important roles in many cutting-edge genomics studies.

## Methods

### The selection and index of seed reads

xRead initially selects P0% of longest reads (default value: 3%) as seed reads at first and assigns zero coverage to all the input reads. Other than the first iteration, xRead selects seed reads with an updated profile of read coverages, i.e., Ps% of the low-covered reads (default value: 10%) are randomly selected as seed reads, where the low-covered reads are defined by a threshold T_RC_ derived from the average coverages of their read parts.

A minimizer-based index is then built for the seed reads. Given a seed read, a set of windows of size W_SR_ (default value: 5bp) starting at every single base are defined and all the *k*-mers (default value: 15bp) within the windows (for both of the strands) are input into a hash function. The k-mer with minimum hash value is chosen to define a quadruple minimizer (*V*_*SR*_, *R*_*SR*_, *P*_*SR*_, *S*_*SR*_ ), where *V*_*SR*_, *R*_*SR*_, *P*_*SR*_ and *S*_*SR*_ indicate the hash value, the read, the position, and the strand of the minimizer. All the minimizers of the seed reads are recorded and sorted by their hash values for indexing, moreover, a hash table of *l*-mers (default value: 11bp) is also built as an auxiliary index data structure to accelerate the retrieval and matching of minimizers in the following step.

### Alignment skeleton-based read overlapping

xRead defines all the reads (all the low covered reads) other than seed reads as query reads in the first iteration (other iterations), and aligns them to seed reads to discover new overlaps to construct (refine) the overlapping graph. The alignment is implemented by the alignment skeleton approach as follows. Given a query read, xRead collects all its minimizers using the same hash function at first. The minimizers are matched to seed reads through the read index and xRead separately merges co-linear matches within the same seed reads to build a set of match blocks (MBs).

xRead uses the MBs as vertices to construct a direct acyclic graph (DAG). Two MBs from the same seed read define an edge if they meet the following conditions:

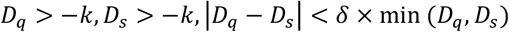

where *D*_*q*_ and *D*_*s*_ are the distances between two MBs on the query and seed reads, respectively, *k* is the maximum allowed overlap length between MBs, and *δ* is a parameter to limit the length difference between *D*_*q*_ and *D*_*s*_. The weight and penalty are also assigned to each edge based on the number of covered bases and the distance between two nodes. The path with the highest score is then inferred in a sparse dynamic programming (SDP) approach by the following recursive equation and is considered the alignment skeleton.

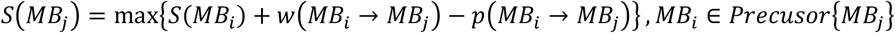

where *S*(*MB*_*j*_) is the score of the vertex *MB*_*j*_, *w*(*MB*_*i*_ →*MB*_*j*_) is the weight of the edge *MB*_*i*_ →*MB*_*j*_, and *p*(*MB*_*i*_ →*MB*_*j*_) is the penalty of the edge *MB*_*i*_ →*MB*_*j*_.

It is also worth noting that multiple alignment skeletons could be built in practice since a query read usually have true positive overlaps to multiple seed reads. More precisely, xRead removes all the MBs along the path after an alignment skeleton is built. Another skeleton is then built with the updated DAG. The iterative process goes on until no alignment skeleton with high score can be built.

### The construction and refinement of core overlapping graph

xRead keeps a global graph data structure to record read overlaps during the iterative process. The produced alignment skeletons are converted to read overlapping information and supplied to the data structure incrementally. For a given query read, the produced alignment skeletons that meet one of the following three conditions are filtered out at first since they could be false positives caused by sequencing errors or repeats in local genomic regions: 1) the overlap length is shorter than T_OM_ (default value: 500bp); 2) the total number of non-redundant bases of all the MBs is higher than T_NB_ (default value: 100bp); 3) the overhang length of either read is longer than T_OH_ (default value: 2000bp). Further, xRead selects at most N_AS_ (default value: 2) highest scored ones of the remaining alignment skeletons as confident read overlaps (CROs) and records their alignment positions to the corresponding two reads.

xRead (re-)estimates read coverages with the updated overlapping information. For a given read, its coverage is estimated by the numbers of the seed reads directly connected to it by the CROs and the reads having CROs to the same seed reads which can be regarded as indirectly aligned to it. It is also worth noting that there could be a proportion of reads being partially overlapped, i.e., some of their read parts have a high number of CROs while other parts have few. Under this circumstance, xRead implements a more precise local estimation, i.e., it splits the given read by W_RC_ size non-overlapping windows (default value: 1000bp) and separately estimates the coverage of various windows by the reads directly and indirectly connected to them. Further, xRead computes the average of the window coverages as the estimated coverage for a read.

The distribution of the coverages of various reads is then estimated and the medium of read coverage M_RC_ is computed. A threshold T_RC_ is set as M_RC_ × *P*_*RC*_ for the selection of seed reads in the next iteration where *P*_*RC*_ is a user-defined parameter (default value: 0.5). With given T_RC_, xRead monitors the number of newly selected seed reads. If it is too low, xRead considers that there are few reads being lowly covered and outputs the resulted graph in PAF format.

### The inference of comprehensive overlapping graph

The graph produced by xRead can be regarded as a core graph consisting of the overlaps between the seed and query reads. As some of the de novo assembly approaches require comprehensive read overlapping information, xRead provides an additional function to infer the overlaps between non-seed reads and produce a more comprehensive graph. Mainly, it implements an iterative width-first searching approach based on the transitive relationships among the overlaps. That is, for each of the non-seed reads, xRead initially retrieves all the other reads connected to the same seed read(s) it attached and infers the overlaps via the transitive relationships. The length and placement of the inferred overlaps are investigated and the ones meeting the conditions similar to that of CROs are remained. Further, the remained overlaps are added to the graph as virtual edges and xRead further expands the graph through them in the following iterations. The transitive-overlap-based inference continues until no new legal overlap is found or it reaches a pre-defined number of iterations.

### The implementation of benchmark

We implemented benchmarks on simulated and real long-read datasets from nine genomes, i.e., *E. coli* (ASM584v2), *S. cerevisiae* (R64), *C. elegans* (WBcel235), *A. thaliana* (TAIR10.1), *D. melanogaster* (Release 6 plus ISO1 MT), *Z. mays* (subsp. *mays* SK) [35], *M. musculus* (GRCm39), *H. sapiens* (GRCh38.p14), *A. mexicanum* (AmbMex60DD) [7]. Most of the datasets are ONT or PacBio CLR reads, and xRead was compared with four state-of-the-art tools, i.e., MHAP (version 2.1.3), MECAT2 (v20190314), Minimap2 (version 2.24) and wtdbg2 (version 2.5). Moreover, a couple of in-house Python scripts were used to interpret and evaluate the outputs of the tools in various formats. Other state-of-the-art assemblers like Canu, Shasta, NECAT, Flye, and Nextdenovo were not included in the benchmarks since they do not provide stand-alone modules to output interpretable results of read overlapping, or employ generic alignment tool such as Minimap2 for read overlapping. All the benchmarks were implemented on a server with 4 Intel Xeon 5220R CPUs (96 CPU cores in total) and 1 Terabyte RAM running Linux Ubuntu 16.04. Refer to Supplementary Tables 3, 4, 7 and 8 (Parameter columns) for the detailed settings of the tools used in the benchmark.

## Declarations

### Ethics approval and consent to participate

Not applicable.

### Consent for publication

Not applicable.

### Availability of data and material

The source code of xRead and the scripts of data simulations are available at: https://github.com/tcKong47/xRead under MIT open source license.

Refer to Additional file 1: Supplementary Notes for the availability of the real sequencing datasets used in benchmark.

### Author contributions

TK implemented the method, BL design the method, and TK, BL and YW performed the analysis. All of the authors wrote the manuscript. TK and BL contributed equally to this work.

## References

1. Eid J, Fehr A, Gray J, Luong K, Lyle J, Otto G, Peluso P, Rank D, Baybayan P, Bettman B, et al: Real-Time DNA Sequencing from Single Polymerase Molecules. Science 2009, 323:133–138.

2. Mikheyev AS, Tin MMY: A first look at the Oxford Nanopore MinION sequencer. Molecular Ecology Resources 2014, 14:1097–1102.

3. Logsdon GA, Vollger MR, Eichler EE: Long-read human genome sequencing and its applications. Nature Reviews Genetics 2020, 21:597–614.

4. Nurk S, Koren S, Rhie A, Rautiainen M, Bzikadze AV, Mikheenko A, Vollger MR, Altemose N, Uralsky L, Gershman A, et al: The complete sequence of a human genome. Science 2022, 376:44-+.

5. Garg S, Fungtammasan A, Carroll A, Chou M, Schmitt A, Zhou X, Mac S, Peluso P, Hatas E, Ghurye J, et al: Chromosome-scale, haplotype-resolved assembly of human genomes. Nature Biotechnology 2021, 39:309–312.

6. Neale DB, Wegrzyn JL, Stevens KA, Zimin AV, Puiu D, Crepeau MW, Cardeno C, Koriabine M, Holtz-Morris AE, Liechty JD, et al: Decoding the massive genome of loblolly pine using haploid DNA and novel assembly strategies. Genome Biology 2014, 15.

7. Nowoshilow S, Schloissnig S, Fei JF, Dahl A, Pang AWC, Pippel M, Winkler S, Hastie AR, Young G, Roscito JG, et al: The axolotl genome and the evolution of key tissue formation regulators. Nature 2018, 554:50-+.

8. Shao C, Sun S, Liu K, Wang J, Li S, Liu Q, Deagle BE, Seim I, Biscontin A, Wang Q, et al: The enormous repetitive Antarctic krill genome reveals environmental adaptations and population insights. Cell 2023.

9. Sovic I, Krizanovic K, Skala K, Sikic M: Evaluation of hybrid and non-hybrid methods for de novo assembly of nanopore reads. Bioinformatics 2016, 32:2582–2589.

10. Jayakumar V, Sakakibara Y: Comprehensive evaluation of non-hybrid genome assembly tools for third-generation PacBio long-read sequence data. Briefings in Bioinformatics 2019, 20:866–876.

11. Rhie A, McCarthy SA, Fedrigo O, Damas J, Formenti G, Koren S, Uliano-Silva M, Chow W, Fungtammasan A, Kim J, et al: Towards complete and error-free genome assemblies of all vertebrate species. Nature 2021, 592:737-+.

12. Lewin HA, Robinson GE, Kress WJ, Baker WJ, Coddington J, Crandall KA, Durbin R, Edwards SV, Forest F, Gilbert MTP, et al: Earth BioGenome Project: Sequencing life for the future of life. Proceedings of the National Academy of Sciences of the United States of America 2018, 115:4325–4333.

13. Berlin K, Koren S, Chin CS, Drake JP, Landolin JM, Phillippy AM: Assembling large genomes with single-molecule sequencing and locality-sensitive hashing. Nature Biotechnology 2015, 33:623-+.

14. Chin CS, Peluso P, Sedlazeck FJ, Nattestad M, Concepcion GT, Clum A, Dunn C, O’Malley R, Figueroa-Balderas R, Morales-Cruz A, et al: Phased diploid genome assembly with single-molecule real-time sequencing. Nature Methods 2016, 13:1050-+.

15. Koren S, Walenz BP, Berlin K, Miller JR, Bergman NH, Phillippy AM: Canu: scalable and accurate long-read assembly via adaptive k-mer weighting and repeat separation. Genome Research 2017, 27:722–736.

16. Xiao CL, Chen Y, Xie SQ, Chen KN, Wang Y, Han Y, Luo F, Xie Z: MECAT : fast mapping, error correction, and de novo assembly for single-molecule sequencing reads. Nature Methods 2017, 14:1072-+.

17. Ruan J, Li H: Fast and accurate long-read assembly with wtdbg2. Nature Methods 2020, 17:155-+.

18. Shafin K, Pesout T, Lorig-Roach R, Haukness M, Olsen HE, Bosworth C, Armstrong J, Tigyi K, Maurer N, Koren S, et al: Nanopore sequencing and the Shasta toolkit enable efficient de novo assembly of eleven human genomes. Nature Biotechnology 2020, 38:1044-+.

19. Chin CS, Alexander DH, Marks P, Klammer AA, Drake J, Heiner C, Clum A, Copeland A, Huddleston J, Eichler EE, et al: Nonhybrid, finished microbial genome assemblies from long-read SMRT sequencing data. Nature Methods 2013, 10:563-+.

20. Chaisson MJ, Tesler G: Mapping single molecule sequencing reads using basic local alignment with successive refinement (BLASR): application and theory. Bmc Bioinformatics 2012, 13.

21. Myers G: Efficient Local Alignment Discovery amongst Noisy Long Reads. In 14th International Workshop on Algorithms in Bioinformatics (WABI); Sep 08-10; Wroclaw, POLAND. 2014: 52–67.

22. Kolmogorov M, Yuan J, Lin Y, Pevzner PA: Assembly of long, error-prone reads using repeat graphs. Nature Biotechnology 2019, 37:540-+.

23. Li H: Minimap2: pairwise alignment for nucleotide sequences. Bioinformatics 2018, 34:3094–3100.

24. Vaser R, Šikić M: Time- and memory-efficient genome assembly with Raven. Nature Computational Science 2021, 1:332–336.

25. Nie F, Huang N, Zhang J, Ni P, Wang Z, Xiao C, Luo F, Wang J: de novo diploid genome assembly using long noisy reads via haplotype-aware error correction and inconsistent overlap identification. bioRxiv 2023:2022.2009.2025.509436.

26. Manavski SA, Valle G: CUDA compatible GPU cards as efficient hardware accelerators for Smith-Waterman sequence alignment. Bmc Bioinformatics 2008, 9.

27. Rognes T: Faster Smith-Waterman database searches with inter-sequence SIMD parallelisation. Bmc Bioinformatics 2011, 12.

28. Daily J: Parasail: SIMD C library for global, semi-global, and local pairwise sequence alignments. Bmc Bioinformatics 2016, 16.

29. Suzuki H, Kasahara M: Introducing difference recurrence relations for faster semi-global alignment of long sequences. Bmc Bioinformatics 2018, 19.

30. Rowe WPM: When the levee breaks: a practical guide to sketching algorithms for processing the flood of genomic data. Genome Biology 2019, 20.

31. Chen Y, Nie F, Xie SQ, Zheng YF, Dai Q, Bray T, Wang YX, Xing JF, Huang ZJ, Wang DP, et al: Efficient assembly of nanopore reads via highly accurate and intact error correction. Nature Communications 2021, 12.

32. Amarasinghe SL, Su S, Dong XY, Zappia L, Ritchie ME, Gouil Q: Opportunities and challenges in long-read sequencing data analysis. Genome Biology 2020, 21.

33. Magi A, Semeraro R, Mingrino A, Giusti B, D’Aurizio R: Nanopore sequencing data analysis: state of the art, applications and challenges. Briefings in Bioinformatics 2018, 19:1256–1272.

34. Carneiro MO, Russ C, Ross MG, Gabriel SB, Nusbaum C, DePristo MA: Pacific biosciences sequencing technology for genotyping and variation discovery in human data. Bmc Genomics 2012, 13.

35. Yang N, Liu J, Gao Q, Gui ST, Chen L, Yang LF, Huang J, Deng TQ, Luo JY, He LJ, et al: Genome assembly of a tropical maize inbred line provides insights into structural variation and crop improvement. Nature Genetics 2019, 51:1052-+.

