## Supplementaly Notes for "xRead: a coverage-guided approach for scalable construction of read overlapping graph"

**Supplementary Notes**

Tangchao Kong^1, 2, 3^, Bo Liu^1, 2, 3, *^, Yadong Wang^1, 2, *^

^1^Center for Bioinformatics, faculty of computing, Harbin Institute of Technology, Harbin, Heilongjiang 150001, China

^2^ Key Laboratory of Biological Bigdata, Ministry of Education, Harbin Institute of Technology, Harbin, Heilongjiang 150001, China

^3^ These authors contributed equally to this work

* Corresponding authors

Contact:

### Supplementary Table 1. Detailed information of reference genomes

| **No.** | **Reference genome** | **Version** | **Genome size** | **Availability** |
| --- | --- | --- | --- | --- |
| 1 | *Escherichia coli* | ASM584v2 | 4.6 Mb | https://www.ncbi.nlm.nih.gov/data-hub/genome/GCF_000005845.2/ |
| 2 | *Saccharomyces cerevisiae* | R64 | 12.1 Mb | https://www.ncbi.nlm.nih.gov/data-hub/genome/GCF_000146045.2/ |
| 3 | *Caenorhabditis elegans* | WBcel235 | 100.3 Mb | https://www.ncbi.nlm.nih.gov/data-hub/genome/GCF_000002985.6/ |
| 4 | *Arabidopsis thaliana* | TAIR10.1 | 119.1 Mb | https://www.ncbi.nlm.nih.gov/data-hub/genome/GCF_000001735.4/ |
| 5 | *Drosophila melanogaster* | Release 6 plus ISO1 MT | 143.7 Mb | https://www.ncbi.nlm.nih.gov/data-hub/genome/GCF_000001215.4/ |
| 7 | *Zea mays (SK)* | GWHAACS00000000 | 2.2 Gb | https://ngdc.cncb.ac.cn/gwh/Assem-bly/123/show |
| 6 | *Mus musculus* | GRCm39 | 2.7 Gb | https://www.ncbi.nlm.nih.gov/data-hub/genome/GCF_000001635.27/ |
| 8 | *Homo sapiens* | GRCh38.p14 | 3.1 Gb | https://www.ncbi.nlm.nih.gov/data-hub/genome/GCF_000001405.40/ |
| 9 | *Ambystoma mexicanum* | AmbMex60DD | 28.2 Gb | https://www.ncbi.nlm.nih.gov/data-hub/genome/GCA_002915635.3/ |

### Supplementary Table 2. Detailed information of simulated datasets

| **No.** | **Reference genome** | **Error model ^a^** | **File size** | **Num. of reads** | **Error rate** |
| --- | --- | --- | --- | --- | --- |
| 1 | *Escherichia coli* | R103 | 443MB | 15378 | 13.0% |
| 2 | *Saccharomyces cerevisiae* | R103 | 1.2GB | 40562 | 13.0% |
| 3 | *Caenorhabditis elegans* | R103 | 9.4GB | 334603 | 13.0% |
| 4 | *Arabidopsis thaliana* | R103 | 12GB | 398748 | 13.0% |
| 5 | *Drosophila melanogaster* | R103 | 14GB | 561293 | 13.0% |
| 6 | *Zea mays (SK)* | R103 | 202GB | 7724943 | 13.0% |
| 7 | *Mus musculus* | R103 | 255GB | 9376380 | 13.0% |
| 8 | *Homo sapiens* | R103 | 291GB | 10390374 | 13.0% |
| 9 | *Ambystoma mexicanum* | R103 | 2TB | 95447945 | 13.0% |

(a) The datasets were simulated by PBSIM2 using the pre-trained R103 chemistry model based on the reference genomes in Table 1. The read-depth, ratio of sequencing errors (mismatches: insertions: deletions), mean length, and total error rate were configured as 50, 23:31:46, 13000, and 13% respectively.

### Supplementary Table 3. The performance of various tools on simulated datasets

| **Tool** | **Parameter ^a^** | **Thread ^b^** | **Real time ^c^** | **CPU time ^c^** | **Memory ^d^** |
| --- | --- | --- | --- | --- | --- |
| ***Escherichia coli*** | | | | | |
| xRead | -k 15 -l 11 -w 5 -x 3 -X 10 -t 8 -M 16 | 8 | 7.962 | 15.273 | 0.348 |
| MHAP | --settings 2 --num_threads 8 | 8 | 51.180 | 440.980 | 16.390 |
| MECAT2 | -outfmt paf -num_threads 8 | 8 | 13.589 | 80.461 | 2.107 |
| minimap2 | -x ava-ont -t 8 | 8 | 18.839 | 112.224 | 3.309 |
| wtdbg2 | -p 0 -k 15 -AS 2 -s 0.05 -L 5000 -t 8 | 8 | 18.243 | 51.800 | 1.418 |
| ***Saccharomyces cerevisiae*** | | | | | |
| xRead | -k 15 -l 11 -w 5 -x 3 -X 10 -t 8 -M 16 | 8 | 19.113 | 47.724 | 0.882 |
| MHAP | --settings 2 --num_threads 8 | 8 | 137.370 | 1213.450 | 20.247 |
| MECAT2 | -outfmt paf -num_threads 8 | 8 | 55.473 | 325.663 | 5.113 |
| minimap2 | -x ava-ont -t 8 | 8 | 53.262 | 342.099 | 5.686 |
| wtdbg2 | -p 0 -k 15 -AS 2 -s 0.05 -L 5000 -t 8 | 8 | 71.104 | 518.350 | 2.705 |
| ***Caenorhabditis elegans*** | | | | | |
| xRead | -k 15 -l 11 -w 5 -x 3 -X 10 -t 8 -M 16 | 8 | 321.739 | 1906.021 | 4.825 |
| MHAP | --settings 2 --num_threads 8 | 8 | 1263.420 | 12322.480 | 77.560 |
| MECAT2 | -outfmt paf -num_threads 8 | 8 | 689.072 | 4681.945 | 34.640 |
| minimap2 | -x ava-ont -t 8 | 8 | 1650.880 | 12039.550 | 21.913 |
| wtdbg2 | -p 0 -k 15 -AS 2 -s 0.05 -L 5000 -t 8 | 8 | 1929.324 | 15271.740 | 18.029 |
| ***Arabidopsis thaliana*** | | | | | |
| xRead | -k 15 -l 11 -w 5 -x 3 -X 10 -t 8 -M 16 | 8 | 329.197 | 1744.481 | 5.586 |
| MHAP | --settings 2 --num_threads 8 | 8 | 3449.579 | 21751.850 | 81.197 |
| MECAT2 | -outfmt paf -num_threads 8 | 8 | 739.716 | 4912.011 | 34.675 |
| minimap2 | -x ava-ont -t 8 | 8 | 1533.620 | 11737.850 | 27.457 |
| wtdbg2 | -p 0 -k 15 -AS 2 -s 0.05 -L 5000 -t 8 | 8 | 2019.105 | 16175.720 | 22.063 |
| ***Drosophila melanogaster*** | | | | | |
| xRead | -k 15 -l 11 -w 5 -x 3 -X 10 -t 8 -M 16 | 8 | 402.298 | 2220.212 | 5.999 |
| MHAP | --settings 2 --num_threads 8 | 8 | 7325.109 | 50070.110 | 98.501 |
| MECAT2 | -outfmt paf -num_threads 8 | 8 | 909.532 | 6077.598 | 34.649 |
| minimap2 | -x ava-ont -t 8 | 8 | 1766.140 | 13441.950 | 37.479 |
| wtdbg2 | -p 0 -k 15 -AS 2 -s 0.05 -L 5000 -t 8 | 8 | 2218.785 | 17815.280 | 26.377 |
| ***Zea mays (SK)*** | | | | | |
| xRead | -k 15 -l 11 -w 5 -x 3 -X 10 -t 24 -M 24 | 24 | 21.93 (h) | 1800879 | 14.018 |
| MHAP | --settings 2 --num_threads 24 | 24 | - | - | - |
| MECAT2 | -outfmt paf -num_threads 24 | 24 | 35.30 (h) | 2950036 | 72.997 |
| minimap2 | -x ava-ont -t 24 | 24 | 277.69 (h) | 5448552 | 64.337 |
| wtdbg2 | -p 19 -AS 2 -s 0.05 -L 5000 -t 24 | 24 | 26.38 (h) | 2133992 | 336.188 |
| ***Mus musculus*** | | | | | |
| xRead | -k 15 -l 11 -w 5 -x 3 -X 10 -t 24 -M 24 | 24 | 12.94 (h) | 878113 | 14.891 |
| MHAP | -settings 2 --num_threads 24 | 24 | - | - | - |
| MECAT2 | -outfmt paf -num_threads 24 | 24 | 31.62 (h) | 2594507 | 73.285 |
| minimap2 | -x ava-ont -t 24 | 24 | 42.07 (h) | 3127297 | 51.953 |
| wtdbg2 | -p 19 -AS 2 -s 0.05 -L 5000 -t 24 | 24 | 35.26 (h) | 2940047 | 292.800 |
| ***Homo sapiens*** | | | | | |
| xRead | -k 15 -l 11 -w 5 -x 3 -X 10 -t 24 -M 24 | 24 | 22.22 (h) | 1583443 | 14.697 |
| MHAP | -settings 2 --num_threads 24 | 24 | - | - | - |
| MECAT2 | -outfmt paf -num_threads 24 | 24 | 50.19 (h) | 4182559 | 73.230 |
| minimap2 | -x ava-ont -t 24 | 24 | 72.97 (h) | 5916894 | 54.097 |
| wtdbg2 | -p 19 -AS 2 -s 0.05 -L 5000 -t 24 | 24 | 50.11 (h) | 4095936 | 326.740 |
| ***Ambystoma mexicanum* ^e^** | | | | | |
| xRead | -k 15 -l 11 -w 5 -x 3 -X 10 -t 64 -M 64 | 64 | 465.16 (h) | 24792.34 (h) | 59.446 |
| MHAP | -settings 2 --num_threads 64 | 64 | - | - | - |
| MECAT2 | -outfmt paf -num_threads 64 | 64 | 1005.52 (h) | 60233.28 (h) | 172.46 |
| minimap2 | -x ava-ont -t 64 | 64 | - | - | - |
| wtdbg2 | -p 19 -AS 2 -s 0.05 -L 5000 -t 64 | 64 | - | - | - |

(a) The parameters of the tools used for benchmarking on the simulated datasets.

(b) The number of CPU threads used for benchmarking on the simulated datasets.

(c) The real-time and CPU time of the tools cost on the simulated datasets, the results marked by “h” indicates CPU hours, otherwise, CPU seconds.

(d) The memory footprints of the tools (in GB) on the simulated datasets.

(e) Only xRead and MECAT2 finished the A. mexicanum dataset. Other tools failed due to out of memory or very large time cost.

‘-’ indicates that the result of the tool is not available for the dataset.

### Supplementary Table 4. The yields of various tools on simulated datasets

| **Tool** | **Parameter ^a^** | **Preci-sion ^b^** | **Sensiti-vity ^b^** | **R% ^c^** | **C% ^d^** | **Gap Num. ^e^** | **Con.**  **Num. ^f^** |
| --- | --- | --- | --- | --- | --- | --- | --- |
|  | ***Escherichia coli*** |  |  |  |  |  |  |
| xRead | -k 15 -l 11 -w 5 -x 3 -X 10 -t 8 -M 16 | 99.939 | 4.339 | 99.668 | 99.989 | 0 | 1 |
| MHAP | --settings 2 --num_threads 8 | 94.437 | 52.282 | 95.981 | 99.989 | 0 | 1 |
| MECAT2 | -outfmt paf -num_threads 8 | 99.760 | 5.697 | 97.022 | 99.989 | 0 | 1 |
| minimap2 | -x ava-ont -t 8 | 97.160 | 94.787 | 99.974 | 99.989 | 0 | 1 |
| wtdbg2 | -p 0 -k 15 -AS 2 -s 0.05 -L 5000 -t 8 | 91.585 | 79.824 | 96.118 | 99.989 | 0 | 1 |
|  | ***Saccharomyces cerevisiae*** |  |  |  |  |  |  |
| xRead | -k 15 -l 11 -w 5 -x 3 -X 10 -t 8 -M 16 | 99.219 | 4.177 | 99.448 | 99.875 | 1 | 2 |
| MHAP | --settings 2 --num_threads 8 | 68.494 | 54.348 | 96.218 | 99.875 | 1 | 1 |
| MECAT2 | -outfmt paf -num_threads 8 | 99.151 | 5.641 | 97.347 | 99.872 | 2 | 4 |
| minimap2 | -x ava-ont -t 8 | 89.861 | 94.681 | 99.958 | 99.875 | 1 | 1 |
| wtdbg2 | -p 0 -k 15 -AS 2 -s 0.05 -L 5000 -t 8 | 54.065 | 78.933 | 96.100 | 99.872 | 2 | 2 |
|  | ***Caenorhabditis elegans*** |  |  |  |  |  |  |
| xRead | -k 15 -l 11 -w 5 -x 3 -X 10 -t 8 -M 16 | 99.709 | 4.295 | 99.645 | 99.993 | 0 | 3 |
| MHAP | --settings 2 --num_threads 8 | 88.509 | 56.490 | 96.599 | 99.993 | 0 | 2 |
| MECAT2 | -outfmt paf -num_threads 8 | 96.017 | 5.865 | 97.812 | 99.993 | 0 | 4 |
| minimap2 | -x ava-ont -t 8 | 46.376 | 95.402 | 99.978 | 99.993 | 0 | 2 |
| wtdbg2 | -p 0 -k 15 -AS 2 -s 0.05 -L 5000 -t 8 | 81.821 | 80.379 | 96.198 | 99.993 | 0 | 2 |
|  | ***Arabidopsis thaliana*** |  |  |  |  |  |  |
| xRead | -k 15 -l 11 -w 5 -x 3 -X 10 -t 8 -M 16 | 99.709 | 4.295 | 99.645 | 99.993 | 0 | 1 |
| MHAP | --settings 2 --num_threads 8 | 88.509 | 56.490 | 96.599 | 99.993 | 0 | 1 |
| MECAT2 | -outfmt paf -num_threads 8 | 96.017 | 5.865 | 97.812 | 99.993 | 0 | 10 |
| minimap2 | -x ava-ont -t 8 | 46.376 | 95.402 | 99.978 | 99.993 | 0 | 1 |
| wtdbg2 | -p 0 -k 15 -AS 2 -s 0.05 -L 5000 -t 8 | 81.821 | 80.379 | 96.198 | 99.993 | 2 | 1 |
|  | ***Drosophila melanogaster*** |  |  |  |  |  |  |
| xRead | -k 15 -l 11 -w 5 -x 3 -X 10 -t 8 -M 16 | 91.126 | 4.095 | 89.684 | 99.993 | 1 | 861 |
| MHAP | --settings 2 --num_threads 8 | 6.585 | 58.543 | 96.567 | 99.994 | 0 | 2 |
| MECAT2 | -outfmt paf -num_threads 8 | 80.760 | 6.498 | 95.006 | 99.993 | 1 | 39 |
| minimap2 | -x ava-ont -t 8 | 22.938 | 90.940 | 96.926 | 99.994 | 0 | 477 |
| wtdbg2 | -p 0 -k 15 -AS 2 -s 0.05 -L 5000 -t 8 | 34.405 | 68.934 | 80.747 | 99.992 | 3 | 9 |
|  | ***Zea mays (SK)*** |  |  |  |  |  |  |
| xRead | -k 15 -l 11 -w 5 -x 3 -X 10 -t 24 -M 24 | 87.828 | 3.091 | 95.855 | 100.0 | 0 | 20 |
| MHAP | -settings 2 --num_threads 24 | - | - | - | - | - | - |
| MECAT2 | -outfmt paf -num_threads 24 | 15.161 | 11.835 | 96.052 | 100.0 | 0 | 5 |
| minimap2 | -x ava-ont -t 24 | 1.516 | 92.374 | 99.876 | 100.0 | 0 | 1 |
| wtdbg2 | -p 19 -AS 2 -s 0.05 -L 5000 -t 24 | 4.446 | 36.434 | 92.978 | 100.0 | 4 | 16 |
|  | ***Mus musculus*** |  |  |  |  |  |  |
| xRead | -k 15 -l 11 -w 5 -x 3 -X 10 -t 24 -M 24 | 94.948 | 3.952 | 96.210 | 100.0 | 2 | 47 |
| MHAP | -settings 2 --num_threads 24 | - | - | - | - | - | - |
| MECAT2 | -outfmt paf -num_threads 24 | 32.779 | 9.072 | 88.857 | 100.0 | 0 | 405 |
| minimap2 | -x ava-ont -t 24 | 4.482 | 90.418 | 99.913 | 100.0 | 0 | 6 |
| wtdbg2 | -p 19 -AS 2 -s 0.05 -L 5000 -t 24 | 44.502 | 68.809 | 94.710 | 100.0 | 1 | 6 |
|  | ***Homo sapiens*** |  |  |  |  |  |  |
| xRead | -k 15 -l 11 -w 5 -x 3 -X 10 -t 24 -M 24 | 92.357 | 4.050 | 95.026 | 100.0 | 0 | 91 |
| MHAP | -settings 2 --num_threads 24 | - | - | - | - | - | - |
| MECAT2 | -outfmt paf -num_threads 24 | 37.482 | 9.521 | 94.372 | 100.0 | 0 | 187 |
| minimap2 | -x ava-ont -t 24 | 6.894 | 90.915 | 99.431 | 100.0 | 0 | 376 |
| wtdbg2 | -p 19 -AS 2 -s 0.05 -L 5000 -t 24 | 59.444 | 71.597 | 94.575 | 100.0 | 0 | 814 |
|  | ***Ambystoma mexicanum* ^g^** |  |  |  |  |  |  |
| xRead | -k 15 -l 11 -w 5 -x 3 -X 10 -t 64 -M 64 | 91.848 | 5.690 | 98.407 | 99.999 | 1 | 862 |
| MHAP | -settings 2 --num_threads 64 | - | - | - | - | - | - |
| MECAT2 | -outfmt paf -num_threads 64 | 4.879 | 26.944 | 97.334 | 99.999 |  |  |
| minimap2 | -x ava-ont -t 64 | - | - | - | - | - | - |
| wtdbg2 | -p 19 -AS 2 -s 0.05 -L 5000 -t 64 | - | - | - | - | - | - |

(a) The parameters of the tools used for benchmarking on the simulated datasets.

(b) The precision and sensitivity of the tools on the simulated datasets.

(c) The proportion of the reads having at least one ground truth overlap being recovered.

(d) The percentage of the donor genome being covered by the connected reads.

(e) The number of gaps on donor genome.

(f) The number of connected components of produced graphs.

(g) Only xRead and MECAT2 finished the A. mexicanum dataset. Other tools failed due to out of memory or very large time cost. Some of the output files of MECAT2 are currently unavailable due to a malfunction of our servers happened recently. The benchmarks are re-implementing and the results will be supplied in the updated version of the manuscript.

‘-’ indicates that the result of the tool is not available for the dataset.

### Supplementary Table 5. The sensitivity of the expanded graphs of xRead on simulated datasets

| **Datasets** | **Outputs of xRead** | | **1 Iteration ^a^** | | **3 Iterations ^a^** | | **5 Iterations ^a^** | |
| --- | --- | --- | --- | --- | --- | --- | --- | --- |
|  | **Sensitivity** | **R%** | **Sensitivity** | **R%** | **Sensitivity** | **R%** | **Sensitivity** | **R%** |
| ***E. coli*** | 4.339 | 99.668 | 87.301 | 99.668 | 98.689 | 99.668 | 98.723 | 99.668 |
| ***S. cerevisiae*** | 4.177 | 99.448 | 86.211 | 99.628 | 98.263 | 99.660 | 98.373 | 99.670 |
| ***C. elegans*** | 4.295 | 99.645 | 85.536 | 99.738 | 98.767 | 99.743 | 98.833 | 99.743 |
| ***A. thaliana*** | 4.287 | 99.472 | 85.841 | 99.681 | 98.365 | 99.690 | 98.468 | 99.691 |
| ***D. melanogaster*** | 4.095 | 89.684 | 77.921 | 95.974 | 93.377 | 96.287 | 93.962 | 96.337 |
| ***Z. mays (SK)*** | 3.091 | 95.855 | 78.712 | 98.037 | 92.887 | 98.794 | 94.472 | 98.922 |
| ***M. musculus*** | 3.952 | 96.210 | 79.968 | 99.238 | 93.608 | 99.412 | 94.903 | 99.490 |
| ***H. sapiens*** | 4.050 | 95.026 | 80.332 | 98.478 | 95.302 | 99.409 | 97.173 | 99.484 |
| ***A. mexicanum* ^b^** | 5.690 | 98.407 | 83.322 | 99.045 |  |  |  |  |

(a) The results of the graphs expanded by 1, 3, and 5 iterations, respectively.

(b) Some output files of xRead on the *A. mexicanum* dataset are currently unavailable due to a malfunction of our servers happened recently. The benchmarks are re-implementing and the results will be supplied in the updated version of the manuscript.

### Supplementary Table 6. Detailed information of real datasets

| **No.** | **Genome** | **Sample** | **Accession** | **Platform** | **Num. of bases** | **Num. of reads** | **Ave. Read length** | **Coverage** |
| --- | --- | --- | --- | --- | --- | --- | --- | --- |
| 1 | *Escherichia coli* | Isolate: EM130d | SRR19746198 | ONT  R93 | 404.6Mb | 34389 | 11766 | 86x |
| 2 | *Caenorhabditis elegans* | strain N2 | SRR10028111 | ONT  R93 | 10.8Gb | 789871 | 13724 | 108x |
| 3 | *Drosophila melanogaster* | BDGP genome strain | SRR13070625 | ONT  R93 | 7.1Gb | 640215 | 11142 | 50x |
| 4 | *Homo sapiens* | HG002 | HG002_ucsc_Oct_2018_Guppy_3.0 | ONT  R93 | 85.6Gb | 10836442 | 7903 | 28x |
| 5 | *Homo sapiens* | HG002 | https://labs.epi2me.io/gm24385_q20_2021.10/ | ONT R10.4 | 224.0Gb | 18125024 | 12360 | 80x |
| 6 | *Homo sapiens* | HG002 | https://ftp-trace.ncbi.nlm.nih.gov/giab/ftp/data/AshkenazimTrio/HG002_NA24385_son | PacBio HiFi | 88.9Gb | 6596012 | 13478 | 28x |
| 7 | *Ambystoma mexicanum* | PRJNA378970 | SRS2051019 | PacBio RS II | 1.28Tb | 105847426 | 9576 | 32x |

### Supplementary Table 7. The performance of various tools on real sequencing datasets

| **Tool** | **Parameter ^a^** | **Thread ^b^** | **Real time ^c^** | **CPU time ^c^** | **Memory ^d^** |
| --- | --- | --- | --- | --- | --- |
| ***Escherichia coli*** | | | | | |
| xRead | -k 15 -l 11 -w 5 -x 3 -X 10 -t 8 -M 16 | 8 | 14.59 | 53.391 | 0.61 |
| MHAP | --setting 2 --num-threads 8 | 8 | 214.92 | 1546.38 | 14.11 |
| MECAT2 | -outfmt paf -num_threads 8 | 8 | 46.58 | 275.91 | 3.28 |
| minimap2 | -x ava-ont -t 8 | 8 | 194.39 | 1171.73 | 5.43 |
| wtdbg2 | -p 0 -k 15 -AS 2 -s 0.05 -L 5000 -t 8 | 8 | 142.33 | 1009.05 | 2.51 |
| ***Caenorhabditis elegans*** | | | | | |
| xRead | -k 15 -l 11 -w 5 -x 3 -X 10 -t 8 -M 16 | 8 | 1531.94 | 10880.96 | 6.08 |
| MHAP | --setting 2 --num-threads 8 | 8 | 22368.01 | 174366.67 | 112.15 |
| MECAT2 | -outfmt paf -num_threads 8 | 8 | 3632.01 | 26397.74 | 30.88 |
| minimap2 | -x ava-ont -t 8 | 8 | 17944.82 | 125949.93 | 47.66 |
| wtdbg2 | -p 0 -k 15 -AS 2 -s 0.05 -L 5000 -t 8 | 8 | 19511.99 | 151825.74 | 36.43 |
| ***Drosophila melanogaster*** | | | | | |
| xRead | -k 15 -l 11 -w 5 -x 3 -X 10 -t 8 -M 16 | 8 | 1324.97 | 9421.72 | 8.22 |
| MHAP | --setting 2 --num-threads 8 | 8 | 29380.11 | 202662.45 | 102.28 |
| MECAT2 | -outfmt paf -num_threads 8 | 8 | 1680.91 | 11798.05 | 35.11 |
| minimap2 | -x ava-ont -t 8 | 8 | 5995.24 | 43179.38 | 39.45 |
| wtdbg2 | -p 0 -k 15 -AS 2 -s 0.05 -L 5000 -t 8 | 8 | 5469.43 | 43568.98 | 25.67 |
| ***Homo sapiens (ONT fast mode)*** | | | | | |
| xRead | -k 15 -l 11 -w 5 -x 5 -X 10 -t 24 -M 24 | 24 | 23.79 (h) | 1652600 | 16.95 |
| MHAP | -settings 2 --num_threads 24 | 24 | - | - | - |
| MECAT2 | -outfmt paf -num_threads 24 | 24 | 20.03 (h) | 1563226 | 76.38 |
| minimap2 | -x ava-ont -t 8 | 24 | 43.90 (h) | 3163418 | 62.74 |
| wtdbg2 | -p 19 -AS 2 -s 0.05 -L 5000 -t 24 | 24 | 26.35 (h) | 2201860 | 157.78 |
| ***Homo sapiens (PacBio HiFi)*** | | | | | |
| xRead | -k 19 -l 11 -w 40 -x 5 -X 10 -t 24 -M 24 | 24 | 2.97 (h) | 48704 | 9.27 |
| MHAP | -settings 2 --num_threads 24 | 24 | - | - | - |
| MECAT2 | -outfmt paf -num_threads 24 | 24 | 27.92 (h) | 2277263 | 68.47 |
| minimap2 | -x ava-pb -t 24 | 24 | 75.76 (h) | 3964617 | 33.21 |
| wtdbg2 | -p 21 -k 0 -AS 4 -K 0.05 -s 0.5 -t 24 | 24 | 12.48 (h) | 1014184 | 110.75 |
| ***Homo sapiens (ONT super high accuracy mode)*** | | | | | |
| xRead | -k 19 -l 11 -w 40 -x 3 -X 10 -t 24 -M 24 | 24 | 14.03 (h) | 386094 | 11.64 |
| MHAP | -settings 2 --num_threads 24 | 24 | - | - | - |
| MECAT2 | -outfmt paf -num_threads 24 | 24 | 134.55 (h) | 11302652 | 70.12 |
| minimap2 | -x ava-ont -t 24 | 24 | 416.37 (h) | 268458775l | 53.47 |
| wtdbg2 | -p 21 -k 0 -AS 4 -K 0.05 -s 0.5 -t 24 | 24 | 42.72 (h) | 3488455 | 316.75 |
| ***Ambystoma mexicanum* ^e^** | | | | | |
| xRead | -k 15 -l 11 -w 5 -x 5 -X 10 -t 64 -M 64 | 64 | 385.92 (h) | 14731.77 (h) | 57.12 |
| MHAP | -settings 2 --num_threads 64 | 64 | - | - | - |
| MECAT2 | -outfmt paf -num_threads 64 | 64 | - | - | - |
| minimap2 | -x ava-ont -t 64 | 64 | - | - | - |
| wtdbg2 | -p 19 -AS 2 -s 0.05 -L 5000 -t 64 | 64 | - | - | - |

(a) The parameters of the tools used for benchmarking on the real datasets.

(b) The number of CPU threads used for benchmarking on the real datasets.

(c) The real-time and CPU time of the tools cost on the real datasets, the results marked by “h” indicates CPU hours, otherwise, CPU seconds.

(d) The memory footprints of the tools (in GB) on the real datasets.

(e) Only xRead finished the *A. mexicanum* dataset. Other tools failed due to out of memory, segmentation fault or very large time cost.

‘-’ indicates that the result of the tool is not available for the dataset.

### Supplementary Table 8. The yields of various tools on real sequencing datasets

| **Tool** | **Parameter ^a^** | **Preci-sion ^b^** | **Sensiti-vity ^b^** | **R% ^c^** | **C% ^d^** | **Gap Num. ^e^** | **Con.**  **Num. ^f^** |
| --- | --- | --- | --- | --- | --- | --- | --- |
|  | ***E. coli*** |  |  |  |  |  |  |
| xRead | -k 15 -l 11 -w 5 -x 3 -X 10 -t 8 -M 16 | 99.795 | 2.935 | 98.810 | 100.0 | 0 | 55 |
| MHAP | --setting 2 --num-threads 8 | 74.929 | 91.140 | 99.907 | 100.0 | 0 | 65 |
| MECAT2 | -outfmt paf -num_threads 8 | 99.811 | 3.706 | 99.764 | 100.0 | 0 | 87 |
| minimap2 | -x ava-ont -t 8 | 93.641 | 96.968 | 99.997 | 100.0 | 0 | 17 |
| wtdbg2 | -p 0 -k 15 -AS 2 -s 0.05 -L 5000 -t 8 | 91.774 | 76.700 | 91.252 | 100.0 | 0 | 60 |
|  | ***C. elegans*** |  |  |  |  |  |  |
| xRead | -k 15 -l 11 -w 5 -x 3 -X 10 -t 8 -M 16 | 99.603 | 2.412 | 98.913 | 100.0 | 0 | 227 |
| MHAP | --setting 2 --num-threads 8 | 57.981 | 85.074 | 98.947 | 100.0 | 0 | 271 |
| MECAT2 | -outfmt paf -num_threads 8 | 99.553 | 5.458 | 98.289 | 100.0 | 0 | 54 |
| minimap2 | -x ava-ont -t 8 | 36.595 | 95.287 | 99.987 | 100.0 | 0 | 636 |
| wtdbg2 | -p 0 -k 15 -AS 2 -s 0.05 -L 5000 -t 8 | 77.976 | 69.450 | 83.782 | 100.0 | 0 | 72 |
|  | ***D. melanogaster*** |  |  |  |  |  |  |
| xRead | -k 15 -l 11 -w 5 -x 3 -X 10 -t 8 -M 16 | 98.183 | 4.217 | 98.226 | 100.0 | 0 | 145 |
| MHAP | --setting 2 --num-threads 8 | 23.767 | 81.457 | 99.922 | 100.0 | 0 | 9 |
| MECAT2 | -outfmt paf -num_threads 8 | 97.482 | 6.816 | 99.012 | 100.0 | 0 | 40 |
| minimap2 | -x ava-ont -t 8 | 23.052 | 78.785 | 99.854 | 100.0 | 0 | 41 |
| wtdbg2 | -p 0 -k 15 -AS 2 -s 0.05 -L 5000 -t 8 | 35.041 | 39.034 | 69.496 | 99.998 | 3 | 11 |
|  | ***H. sapiens (ONT fast mode)*** |  |  |  |  |  |  |
| xRead | -k 15 -l 11 -w 5 -x 5 -X 10 -t 24 -M 24 | 99.264 | 7.214 | 97.642 | 99.995 | 20 | 2530 |
| MHAP | -settings 2 --num_threads 24 | - | - | - | - | - | - |
| MECAT2 | -outfmt paf -num_threads 24 | 90.847 | 55.528 | 98.820 | 99.991 | 26 | 1182 |
| minimap2 | -x ava-ont -t 8 | 7.013 | 90.565 | 99.728 | 100.0 | 2 | 471 |
| wtdbg2 | -p 19 -AS 2 -s 0.05 -L 5000 -t 24 | 54.880 | 26.899 | 43.057 | 99.974 | 197 | 44 |
|  | ***H. sapiens (PacBio HiFi)*** |  |  |  |  |  |  |
| xRead | -k 19 -l 11 -w 40 -x 5 -X 10 -t 24 -M 24 | 99.061 | 7.426 | 99.589 | 99.999 | 3 | 58 |
| MHAP | -settings 2 --num_threads 24 | - | - | - |  | - | - |
| MECAT2 | -outfmt paf -num_threads 24 | 85.673 | 70.471 | 99.982 | 100.0 | 0 | 1 |
| minimap2 | -x ava-pb -t 24 | 1.703 | 95.709 | 99.988 | 100.0 | 0 | 2 |
| wtdbg2 | -p 21 -k 0 -AS 4 -K 0.05 -s 0.5 -t 24 | 96.208 | 77.871 | 99.885 | 99.999 | 2 | 861 |
|  | ***H. sapiens (ONT super high accuracy mode)*** | |  |  |  |  |  |
| xRead | -k 19 -l 11 -w 40 -x 3 -X 10 -t 24 -M 24 | 97.532 | 3.085 | 97.887 | 99.996 | 72 | 2248 |
| MHAP | -settings 2 --num_threads 24 | - | - | - | - | - | - |
| MECAT2 | -outfmt paf -num_threads 24 | 85.836 | 65.911 | 99.955 | 100.0 | 1 | 120 |
| minimap2 | -x ava-ont -t 24 | 1.353 | 94.067 | 99.850 | 99.999 | 26 | 269 |
| wtdbg2 | -p 21 -k 0 -AS 4 -K 0.05 -s 0.5 -t 24 | 95.093 | 64.328 | 89.841 | 99.979 | 161 | 1450 |
|  | ***A. mexicanum ^g^*** |  |  |  |  |  |  |
| xRead | -k 15 -l 11 -w 5 -x 5 -X 10 -t 64 -M 64 | 86.286 | 2.193 | 92.080 |  |  |  |
| MHAP | -settings 2 --num_threads 64 | - | - | - | - | - | - |
| MECAT2 | -outfmt paf -num_threads 64 | - | - | - | - | - | - |
| minimap2 | -x ava-ont -t 64 | - | - | - | - | - | - |
| wtdbg2 | -p 19 -AS 2 -s 0.05 -L 5000 -t 64 | - | - | - | - | - | - |

(a) The parameters of the tools used for benchmarking on the simulated datasets.

(b) The precision and sensitivity of the tools on the simulated datasets.

(c) The proportion of the reads having at least one ground truth overlap being recovered.

(d) The percentage of the donor genome being covered by the connected reads.

(e) The number of gaps on donor genome.

(f) The number of connected components of produced graphs.

(g) Only xRead finished the *A. mexicanum* dataset. Other tools failed due to out of memory, segmentation fault or very large time cost. Some of the output files of xRead are currently unavailable due to a malfunction of our servers happened recently. The benchmarks are re-implementing and the results will be supplied in the updated version of the manuscript.

‘-’ indicates that the result of the tool is not available for the dataset.

### Supplementary Table 9. Percentages of the reads not correctly overlapped by xRead with various causes

| **Datasets** | **R% of xRead** | **R_1_% ^d^** | **R_2_% ^e^** | **R_3_% ^f^** |
| --- | --- | --- | --- | --- |
| ***E. coli*** | 98.810 | 0.000 | 0.028 | 1.162 |
| ***C. elegans*** | 98.913 | 0.014 | 0.052 | 1.021 |
| ***D. melanogaster*** | 98.226 | 0.040 | 1.004 | 0.730 |
| ***H. sapiens ^a^*** | 97.642 | 0.077 | 0.474 | 1.807 |
| ***H. sapiens ^b^*** | 99.589 | 0.179 | 0.193 | 0.039 |
| ***H. sapiens ^c^*** | 97.887 | 0.354 | 0.788 | 0.971 |
| ***A. mexicanum ^g^*** | 92.080 |  |  |  |

(a) The human datasets in ONT fast base-calling modes.

(b) The PacBio HiFi human dataset.

(c) The human datasets in ONT super high accuracy base-calling modes.

(d) The percentage of the reads having not been overlapped.

(e) The percentage of the reads having only false positive overlaps.

(f) The percentage of the reads being overlapped with other reads not included in the pseudo-ground truth set.

(g) Some output files of xRead on the *A. mexicanum* dataset are currently unavailable due to a malfunction of our servers happened recently. The benchmarks are re-implementing and the results will be supplied in the updated version of the manuscript.

### Supplementary Table 10. The sensitivity of the expanded graphs of xRead on real sequencing datasets

| **Datasets** | **Outputs of xRead** | | **1 Iteration ^d^** | | **3 Iterations ^d^** | | **5 Iterations ^d^** | |
| --- | --- | --- | --- | --- | --- | --- | --- | --- |
|  | **Sensitivity** | **R%** | **Sensitivity** | **R%** | **Sensitivity** | **R%** | **Sensitivity** | **R%** |
| ***E. coli*** | 2.935 | 98.810 | 78.177 | 99.015 | 95.625 | 99.018 | 96.067 | 99.018 |
| ***C. elegans*** | 2.412 | 98.913 | 65.082 | 99.039 | 95.284 | 99.052 | 96.360 | 99.056 |
| ***D. melanogaster*** | 4.217 | 98.226 | 71.773 | 98.956 | 95.833 | 99.202 | 96.799 | 99.301 |
| ***H. sapiens ^a^*** | 7.214 | 97.642 | 71.776 | 98.114 | 88.651 | 98.192 | 92.293 | 98.193 |
| ***H. sapiens ^b^*** | 7.426 | 99.589 | 68.439 | 99.915 | 87.833 | 99.926 | 93.073 | 99.929 |
| ***H. sapiens ^c^*** | 3.085 | 97.887 | 49.947 | 98.480 | 86.241 | 98.542 | 88.710 | 98.554 |
| ***A. mexicanum ^e^*** | 2.193 | 92.080 |  |  |  |  |  |  |

(a) The human datasets in ONT fast base-calling modes.

(b) The PacBio HiFi human dataset.

(c) The human datasets in ONT super high accuracy base-calling modes.

(d) The results of the graphs expanded by 1, 3, and 5 iterations, respectively.

(e) Some output files of xRead on the *A. mexicanum* dataset are currently unavailable due to a malfunction of our servers happened recently. The benchmarks are re-implementing and the results will be supplied in the updated version of the manuscript.
